# scRCA: a Siamese network-based pipeline for the annotation of cell types using imperfect single-cell RNA-seq reference data

**DOI:** 10.1101/2024.04.08.588510

**Authors:** Yan Liu, Chen Li, Long-Chen Shen, He Yan, Guo Wei, Robin B. Gasser, Xiaohua Hu, Jiangning Song, Dong-Jun Yu

## Abstract

A critical step in the analysis of single-cell transcriptomic (scRNA-seq) data is the accurate identification and annotation of cell types. Such annotation is usually conducted by comparative analysis with known (reference) data sets – which assumes an accurate representation of cell types within the reference sample. However, this assumption is often incorrect, because factors, such as human errors in the laboratory or in silico, and methodological limitations, can ultimately lead to annotation errors in a reference dataset. As current pipelines for single-cell transcriptomic analysis do not adequately consider this challenge, there is a major demand for a computational pipeline that achieves high-quality cell type annotation using imperfect reference datasets that contain inherent errors (often referred to as “noise”). Here, we built a Siamese network-based pipeline, termed scRCA, that achieves an accurate annotation of cell types employing imperfect reference data. For researchers to decide whether to trust the scRCA annotations, an interpreter was developed to explore the factors on which the scRCA model makes its predictions. We also implemented 3 noise-robust losses-based cell type methods to improve the accuracy using imperfect dataset. Benchmarking experiments showed that scRCA outperforms the proposed noise-robust loss-based methods and methods commonly in use for cell type annotation using imperfect reference data. Importantly, we demonstrate that scRCA can overcome batch effects induced by distinctive single cell RNA-seq techniques. We anticipate that scRCA (https://github.com/LMC0705/scRCA) will serve as a practical tool for the annotation of cell types, employing a reference dataset-based approach.

## Introduction

Single-cell sequencing (scRNA-seq) has emerged as a powerful approach to explore gene transcription/expression (i.e., transcriptomics) at a cellular resolution [1], allowing for in-depth molecular studies using healthy and diseased tissues [2-4]. The accurate identification and annotation of cell types is a critical step in the analysis of single-cell sequence data and serves to uncover cellular heterogeneity within and among tissues, developmental stages, and organisms [5]. However, it can be challenging to accurately annotate cell types; manual annotation can be time-consuming, labor-intensive, and highly dependent on defining marker genes [6, 7]. Recently, numerous annotation methods have been developed which rely on the use of reference datasets – i.e., well-annotated scRNA-seq data [8-16]. Basically, these methods were trained with such reference datasets to define numerous classifiers for subsequent cell type annotation in test datasets, rather than employing known cell type-specific marker genes [17, 18]. The key prerequisite for reference dataset-trained methods, though, is that the cell types have been accurately and reliably annotated in the reference dataset. However, in practice, many factors (e.g., human errors during experimentation, cell or tissue contamination, and/or methodological limitations) can lead to annotation errors (‘noise’) in the reference datasets, which can adversely impact subsequent analyses of test samples and the final quality of annotation.

Recently, much effort has focused on developing methods that learn accurate feature representations from noisy data, particularly in the field of computer vision [19]. In addition to the commonly-used categorical cross-entropy (CCE) loss, many noise-robust loss functions have been proposed, including the forward loss (FW) [20], generalized cross-entropy loss (GCE) [21] and determinant-based mutual information (DMI) [22]. However, single-cell data are distinct in nature from image data - they are usually relatively sparse and contain many false zero-values [23, 24]. Therefore, it is critical to explore the capability of these loss functions to reduce the impact of noise in reference datasets. Moreover, although some cell type annotation algorithms based on deep learning have attained encouraging performance [25, 26], they represent ‘black boxes’ to most biologists [27, 28], because there is little insight to understand how the internal models of these algorithms co-operatively work. Thus, there is a need to address the above issues by developing an algorithm that accurately assigns cell types and provides clear reasoning for the annotations.

Here, we built a reliable and robust pipeline, termed scRCA, by integrating a Siamese network [29] and CCE loss functions, to learn cell type-specific information from reference datasets with errors (‘noise’) that are intrinsic to single-cell studies. To the best of our knowledge, scRCA is the first deep-learning-based computational pipeline which is dedicated to cell type annotation using reference datasets containing noise. To improve the model’s interpretability, we designed and implemented an ‘interpreter’, which defines marker genes required to classify cell types. The results of our experiments showed that scRCA outperforms other state-of-the-art methods for the annotation of cell types, even using suboptimal reference datasets. Importantly, we show that, regardless of RNA-seq technique, scRCA can overcome batch effects and accurately annotate cell types. Thus, scRCA, equipped with an interpreter, should serve as a practical pipeline for the accurate annotation of cell types, employing imperfect reference datasets.

## Materials and Methods

### Benchmark dataset collection and quality control

In this study, a total of 13 PbmcBench datasets [30] were employed to evaluate and benchmark all methods. These datasets were generated using different sequencing protocols, including inDrop (iD), CEL-Seq2 (CL), SMART-Seq2 (SM2), 10X version 2 (10Xv2), 10X version 3 (10Xv3), Seq-Well (SW), and Drop-Seq (DR). All PbmcBench datasets were obtained from the Broad Institute Single Cell portal (https://singlecell.broadinstitute.org/single_cell/study/SCP424/single-cellcomparison-pbmc-data). Following Abdelaal et al. [30] , we then performed a three-step preprocessing to further improve the data quality, including (i) eliminating cells with controversial labels (i.e., doublets, debris, or unlabeled cells); (ii) removing the genes with zero counts across all cells and calculating the median numbers of the detected genes in each cell; and (iii) calculating the median absolute deviation (MAD) across all cells and removing cells when the total number of detected genes was below three MAD values from the median number of detected genes in each cell. The raw data was then normalized via the log normalization method. The pre-processed datasets were described in **Supplementary Table S1**.

### Simulating the label noise

To build predictive models, we synthetically corrupted these well-annotated datasets utilizing the approach proposed by Reed *et al* [31]. We applied the noise transition matrix *M* to these datasets for adding label noise:

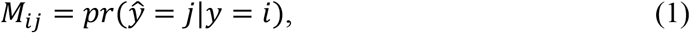

where *y* and *ŷ* represent the correct and flipped annotations for the cell respectively, and *pr*() denotes the probability value. Here, two types of noise transition matrices are used in this study, including pair flip (P) and symmetric flip (S) (**Fig. 1A**). The pair flip structure simulates the scenario where a labeled class can be flipped to the adjacent class instead of a far-away class, while the symmetric flip structure can uniformly flip the labeled class into other classes. In this study, we used S(P)-L% to represent the noise type and level are symmetric flip (pair flip) and L%.

**Fig. 1.**
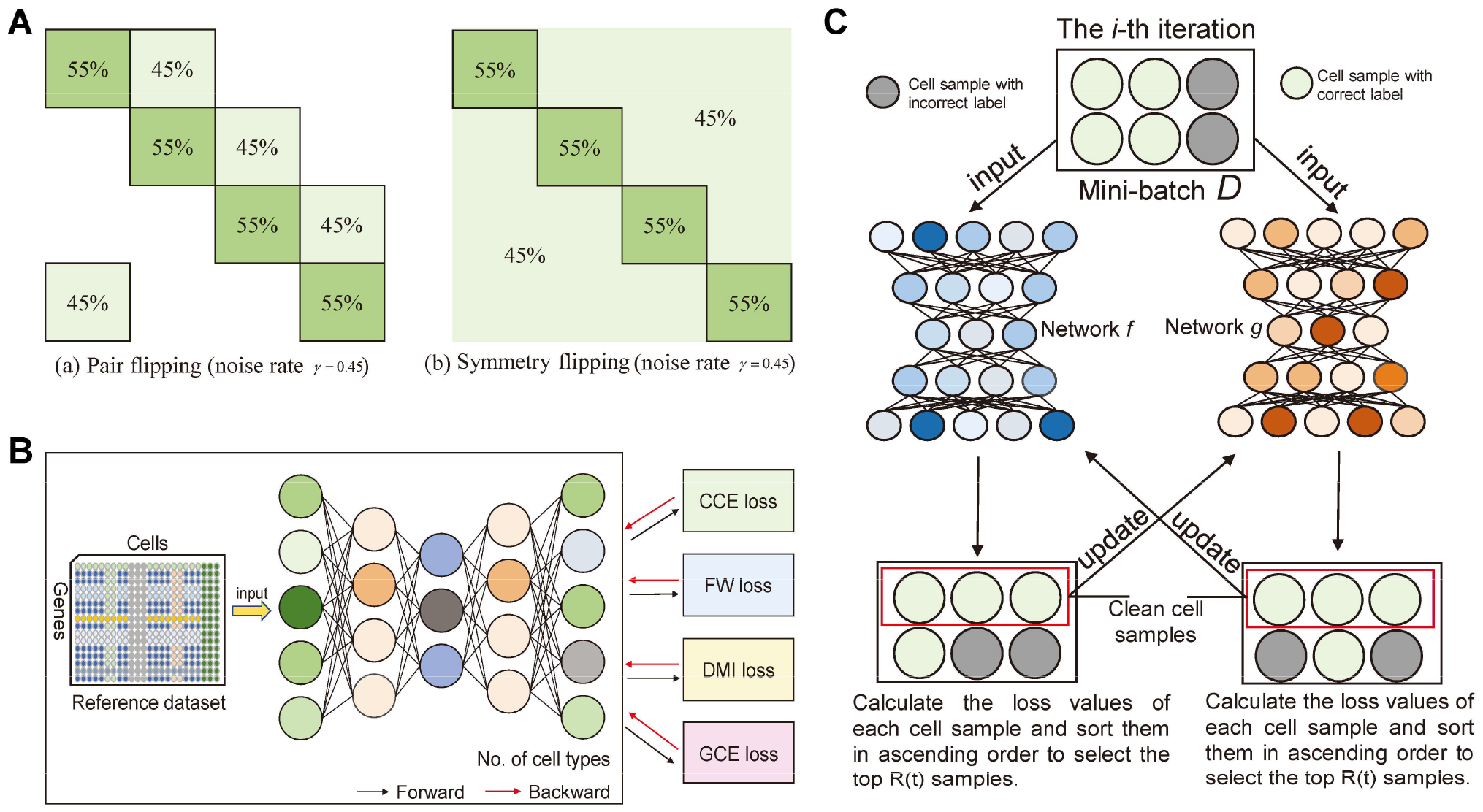
Details of various cell type annotation methods. (A) Transition matrices of different noise types (using four classes as an example); (B) Overview of several benchmark cell type annotation methods, which are based on the noise-robust loss function. The black box is the shared part of each benchmark method; the other methods, constructed with each loss, are independent from one another (CCE loss, FW loss, GCE loss and DMI loss were used to construct scCCE, scFW, scGCE and scDMI respectively); (C) The i-th iteration in the training process.

## Experimental scenarios

To effectively evaluate the cell type annotation across different protocols, amongst the 13 Pbmcbench datasets, we used the pbmc1 dataset of one protocol as the reference dataset (i.e., the training protocol), while the Pbmc1 datasets sequenced by other protocols and the Pbmc2 datasets of the training protocol were combined as the query dataset [32, 33]. We then corrupted the cell types in the reference dataset using the strategy of pair flip and symmetric flip (with the noise rate ranging from 0.15 to 0.45 with a step of 0.1), respectively. The cell types of the query dataset remained unchanged for performance evaluation.

### The scRCA pipeline

In scRCA, we employed a Siamese deep neural network, which has been successfully applied in the field of computer vision [34-37], to co-learn the potential relationships between gene expression profiles and cell types. As shown in **Fig. 1C**, in each mini-batch dataset, each subnetwork selects its small-loss instances as useful knowledge and teaches the knowledge to the other network for training purposes. Specifically, scRCA was constructed to reflect the two important properties of deep networks: (1) the samples with small-loss projected by deep networks after preliminary training are easy to learn - they are more likely to belong to clean samples than large-loss samples [38]; (2) the ‘memorization’ effect of deep networks [38, 39] - despite the incorrect annotations in the reference dataset, scRCA still learns the clean and easy patterns in the initial epochs. Therefore, in the scRCA framework, we employed two networks *f* and *g* to eliminate the noisy samples using their loss values. The calculation details of scRCA are described in **Algorithm 1**.

#### Algorithm 1: scRCA

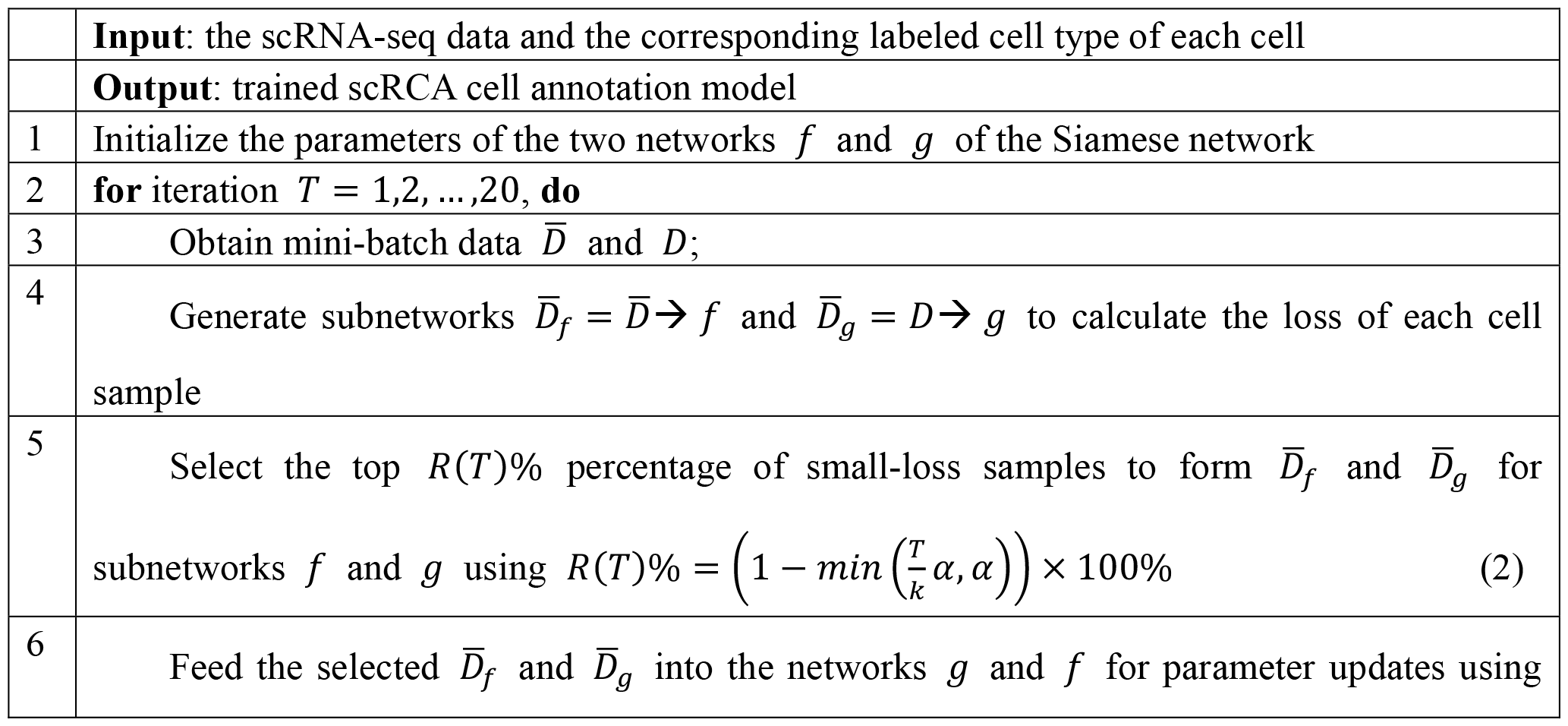
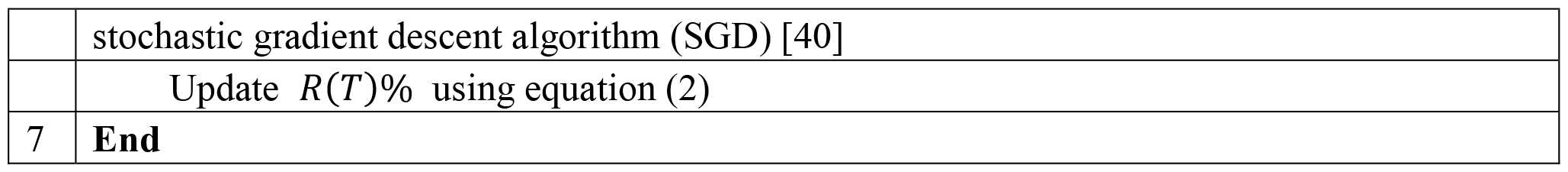

In this study, networks *f* and *g* share the same structure – i.e., each network contains an input layer, two hidden layers, and an output layer. The number of nodes in the input layer nodes is equal to the number of common genes shared between the reference and query datasets. The two hidden layers have 128 and 64 nodes, respectively. The number of nodes in the output layer is equal to the number of cell types in the reference data. The cross-entropy function was used as the loss function of the networks *f* and *g*.

### The interpretation module in scRCA

We applied the loss function proposed by Ribeiro *et al* [41] to construct the interpretation module in scRCA:

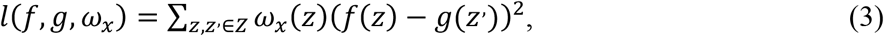

where *f*() and *g*() are the prediction model (i.e., the network *g*) and the interpretation model (i.e., a linear model) of scRCA, respectively. Assuming that x ∈ *R*^*d*^ is the original gene expression of cell sample *and x*′ ∈ {0,1}^*d*^ denotes its interpretable binary representation of presence/absence of a gene, to learn the parameters of interpretation model, we need to sample the cell around *x*′ by drawing nonzero elements of *x*′ randomly, namely perturbed interpretable sample *z*′ ∈ {0,1}^*d*^ . Then, we recover the perturbed sample z ∈ *R*^*d*^ using the perturbed interpretable sample. Specifically, when the value of a given gene is 0 in *z*′, we assign a value of 0 to that gene in z. Otherwise, we assign the value of that gene in x to the value of that gene in z. Finally, we repeat this sampling process 5000 times to learn the parameters of *g*(). *ω*_*x*_ measures the similarity between the cell *z* to *x*:

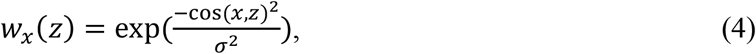

where cos(*A, B*) is the cosine distance of vectors *A* and *B*, σ is the hyperparameter of the RBF kernel to define the characteristic length-scale for learning the similarity between samples, i.e., the ratio of the distance between samples before and after the feature space mapping in the weight space perspective. In this paper, we used the least squares [42] to learn the parameters of liner model, which can be seen as the corresponding contribution value of genes.

### Assessing the performance of various noise-robust loss functions in cell type annotation

scRCA employs CCE as the loss function. We also employed other four widely used loss functions, including CCE (categorical cross-entropy) loss, FW (forward) loss [20], DMI (determinant-based mutual information) loss [22], and generalized cross-entropy loss (GCE) loss [21], to implement four benchmarking methods of scRNA (**Fig. 1B**) for cell type annotation, namely scCCE, scFW, scDMI, and scGCE. Compared to scRCA, these methods have only one network (i.e., network *f*). Refer to **Supplementary Texts S1-3** for a detailed description of these loss functions.

### Identifying potential marker genes for different cell types

The interpretation model of scRCA allows us to identify the genes used by the prediction module of scRCA for cell type annotation, which are termed determining genes. Herein, we further explored the determining genes of scRCA to obtain the genes with expression trends in the query cells, thereby further identifying the potential marker genes of the corresponding cell. To obtain the potential marker genes for different cell types, we first counted the potential marker genes of query cells that are correctly predicted by the scRCA model with a probability greater than 0.99. Second, we selected the 30 genes with the highest frequency of occurrence as the potential marker genes for this cell type.

### Measuring the performance of scRCA

Given that cell type annotation is a multiclass prediction task, we applied several widely used measures for multiclass classification, including weighted-F1score and the overall accuracy, to evaluate the overall performance of our proposed methods. For the weighted-F1 score, first, we calculated the Precision and Recall of each cell type as follows:

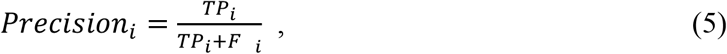

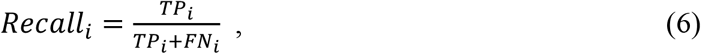

where TP, FP, FN are the true positive, false positive and False Negative, respectively. We then calculated the F1 score of each cell type and the weighted-F1 score as follows:

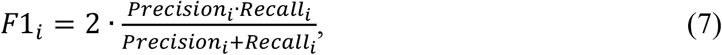

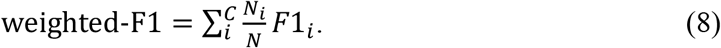

The overall accuracy refers to the proportion of cell samples that are correctly predicted among the total number of query cells:

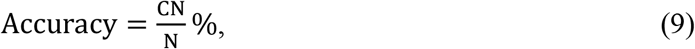

where CN refers to the number of correctly classified query cells, and N denotes to the number of the query cells.

## RESULTS

### Overview of scRCA

Compared with state-of-the-art reference dataset-based cell type annotation methods, scRCA focuses on: (i) enabling cell type annotation based on imperfect reference datasets, and (ii) facilitating the interpretation of cell type annotation results. In practice, it is inevitable that some errors in annotation are introduced – through computational and/or experimentational procedures. Thus, designing a method for the accurate annotation of cell types using imperfect reference datasets will have major benefits, particularly if it can define cell type-specific marker genes [43] to enable the study of biological processes and cellular functions [44]. In this context, we constructed scRCA to include two main steps: (i) model learning and (ii) model interpreter construction (**Fig. 2** and ‘Materials and Methods’). At the model learning stage (**Fig. 2A**), the gene transcription matrix and the corresponding cell types are used as inputs into scRCA. Then, the relationships between cell types and genes are learned and processed in the Siamese network (i.e., *g* and *f*) using CCE as the loss function. At the cell type annotation stage (**Fig. 2B**), we fixed the parameter of the networks *f* and *g*, and then fed the gene transcription matrix of the query cells into the network *g*. Subsequently, the network *g* provides the probability for each query cell representing a particular reference cell type, and the interpretation model of scRCA outputs the gene(s) that the model uses to predict individual query cells.

**Fig. 2.**
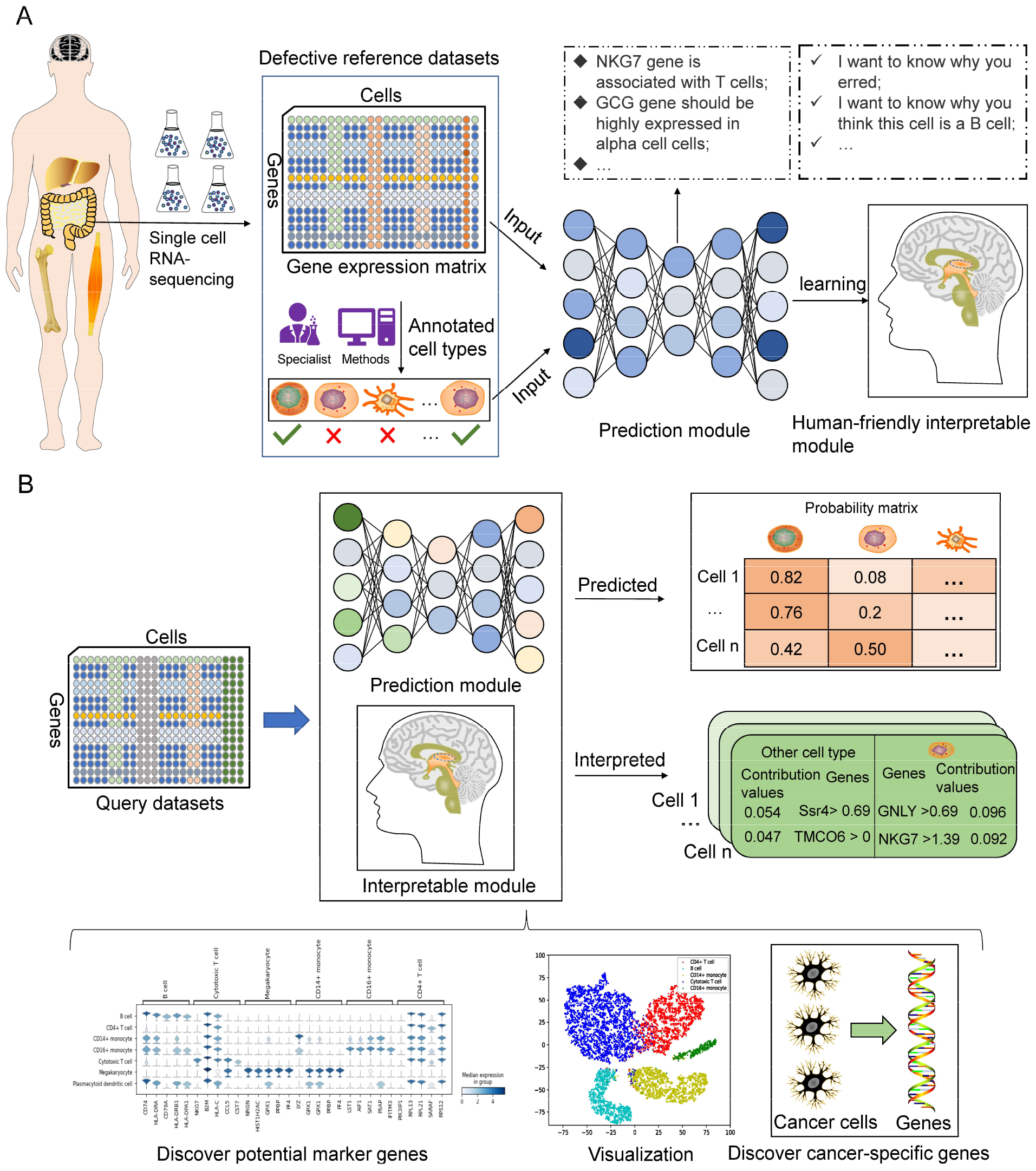
scRCA pipeline. (A) Model learning: the gene transcription matrix and the corresponding cell type are fed into scRCA to learn the relationship between cell types and genes. (cells marked with a red ‘X’ were incorrectly annotated). (B) Annotation: cell types are assigned, and marker genes are identified. Cell type-specific genes enable the discovery of marker gene candidates, visualization, and the discovery of cancer-specific genes.

### CCE outperforms other loss functions in cell type annotation on noisy reference datasets

We evaluated the prediction performance of scRCA (using CCE as the loss function) with different noise-robust loss functions-based cell type annotation methods, including single cell (sc)CCE, scGCE, scFW, and scDMI. In the model training process, we used different training strategies and parameters. First, we employed the fixed parameters to train these models by setting the batch size, learning rate, and epoch to 256, 0.01, and 20, respectively. We employed various noise levels and corruption strategies to corrupt reference datasets. We did not separate a validation dataset from the reference data to identify the appropriate parameters for each network, due to the annotation errors in the reference data. According to **Fig. 3(A-B)**, most cell type annotation methods performed poorly, except scRCA. Specifically, the accuracy of scCCE, scFW, scDMI, and scGCE remained < 40%, regardless of the corruption strategy or noise level. On the contrary, scRCA performed consistently well under all experimental conditions. We assume that the possible reason is that different neural network structures require different hyperparameters for fitting, e.g., learning rate, batch size, and epoch.

**Fig. 3.**
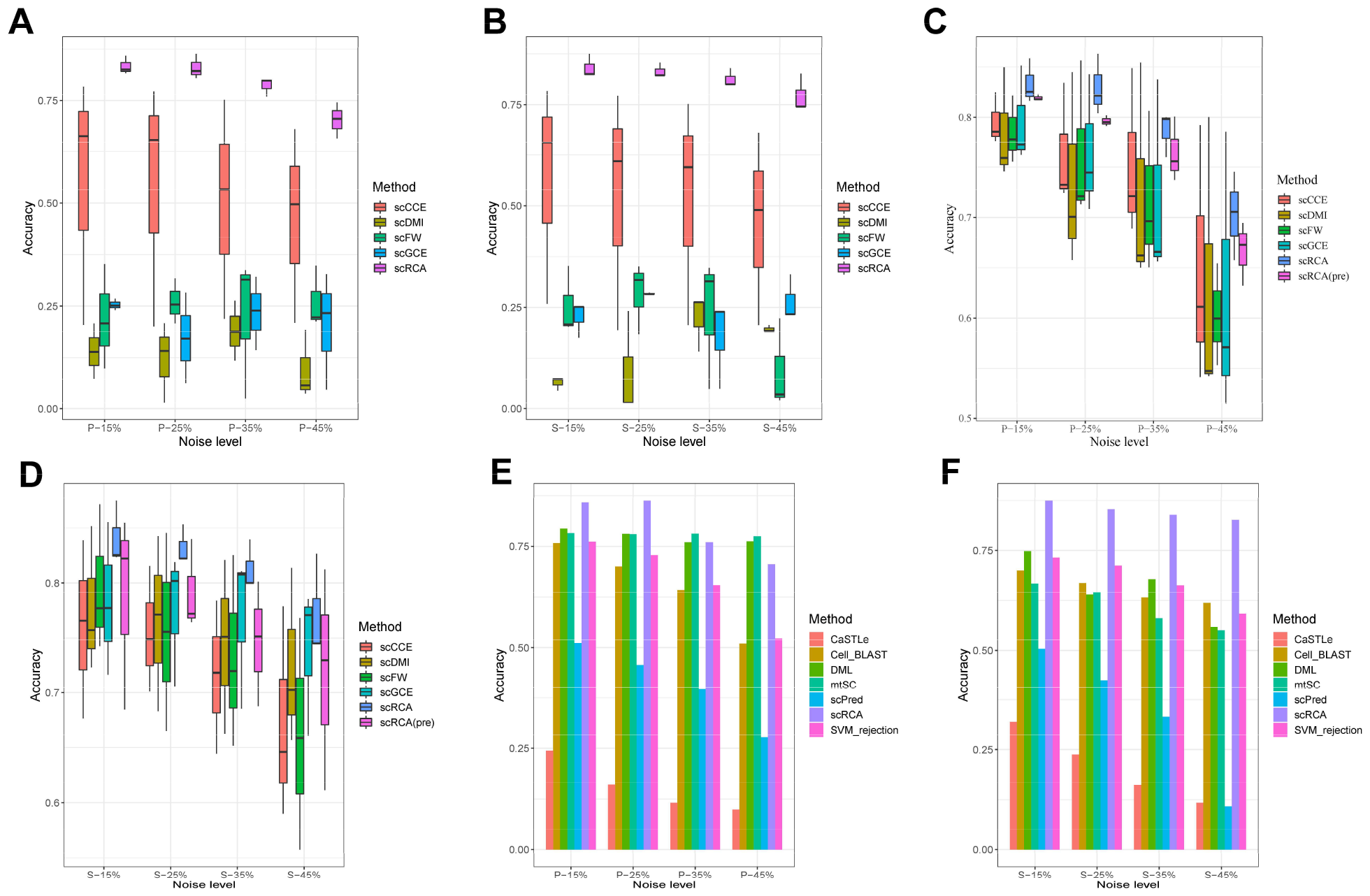
(A-B) The accuracy of cell type annotation for individual methods under different noise types employing the same training strategy (pbmc1_10Xv2, pbmc1_ID, and pbmc1_10Xv3 are the reference dataset respectively); (C-D) The accuracy of cell type annotation for individual methods under different noise types employing the distinct training strategy (pbmc1_10Xv2, pbmc1_ID, and pbmc1_10Xv3 are the reference dataset respectively); (E-F) Benchmarking of scRCA against available methods of cell type annotation employing distinct annotation corruption strategies and levels (pbmc1_ID dataset is the reference dataset).

We then applied different strategies and parameters to train and optimize each model. Specifically, for scCCE, the batch size and epoch were set at 256 and 120, respectively. We used a stepwise strategy to adjust the learning rate, which was set at 0.01, 0.001, and 0.0001 using epochs of 1-40, 40-80, and 80-120, respectively. For scFW, scDMI, scGCE, and scRCA, we first used the CCE loss function to pre-train the network. Then, the batch sizes, learning rates, and epochs of scFW and scDMI were set, in accordance with scCCE. We found that scGCE was sensitive to the batch size, learning rate, and epoch. Therefore, we empirically set the batch size, learning rate, and epoch of scGCE at 16, 0.001, and 400, respectively; the batch size, learning rate, and epoch for scRCA were set at 256, 0.01, and 20, respectively. The performance of all methods was significantly improved (**Fig. 3(C-D)** and **Supplementary Fig. S1**; weighted-F1), except for scRCA. However, the accuracy of scRCA, built using the pre-training strategy, did not improve, which might be explained by the scRCA model using CCE as the loss function of networks *f* and *g*. However, the pre-training process directly uses the entire reference dataset as the training set without eliminating erroneous data (‘noise’). As shown in **Fig. 3(C-D)**, scRCA outperformed all the other methods using different corruption strategies and noise levels. These results (**Fig. 3(C-D)**) indicate that scRCA provides a more accurate annotation of cell types than other noise-robust loss functions-based cell type annotation methods when the reference dataset is imperfect.

### Comparison of the performance of scRCA with other reference dataset-based cell type annotation methods

We evaluated and compared the performance of scRCA with state-of-the-art reference dataset-based cell type annotation methods, including CaSTLe [45], scPred [46], SVM_rejection [47], Cell_Blast [48], mtSC [49], and DML [49]. Similarly, the reference datasets were corrupted using pair-flip and symmetric-flip strategies with various noise levels. According to **Fig. 3(E-F)** and **Supplementary Fig. S2**, regardless of the dataset corruption strategy, the prediction performance of all methods decreased with increasing noise introduced into the reference dataset (**Fig. 3(E-F)**). However, scRCA outperformed other methods, except when the reference dataset was corrupted by pair-flip with a noise level of 45%. DML and mtSC are metric-learning methods that only consider whether the cell types are the same, and do not take specific cell types into account when constructing training samples. As a result, the accuracy of cell type annotation is stable as the noise rate increases when pairwise flipping was performed on the cell types of the reference dataset. Specifically, using the reference dataset, corrupted by symmetric flip and with a noise level of 15%, scRCA achieved an accuracy of ∼87%, which was 12% higher than DML, the second-best method. Furthermore, based on the noise level in the reference datasets (15%–45%) via symmetric flip, scRCA consistently maintained a satisfactory performance. Taken together, using imperfect reference datasets, scRCA could still achieve a sound performance for cell type annotation. These results also demonstrate that in practical applications, when the quality of the reference dataset is unknown, scRCA can still deliver reliable and robust cell annotation predictions.

### Cell type features enable accurate annotation by scRCA

We investigated the properties of the cell features calculated by scRCA for cell type annotation. An ideal feature set should be cell type-specific – i.e., identical cell types share the same or similar features, while cells of different types have distinct features. Therefore, we ran bi-clustering to analyze the features generated by scRCA for different cell types. Specifically, we randomly used the corrupted pbmc1_ID with S-15% (symmetric flip; 15% of noise level) dataset as the reference dataset to train the networks *f* and *g*. We fed the gene transcription matrix of the query dataset into the trained network *g* and then collected the outputs of the second hidden layer as the cell type (CT) features (64 dimensions). To objectively analyze the features extracted, we generated two datasets with 100 cells each, randomly selected from pbmc1_10Xv2 dataset and the combined query dataset, respectively. Specifically, the first 100 cells were selected randomly from the pbmc1_10Xv2 dataset representing five cell types; the second 100 cells (covering 7 cell types) were then randomly selected from the combined query dataset, which includes all other pbmc1datasets and the pmbc2_ID dataset. Finally, we performed a bi-clustering analysis using the CT features and the raw gene transcription data for the selected cells (**Fig. 4A, Supplementary Figs. S3-S5**).

**Fig. 4.**
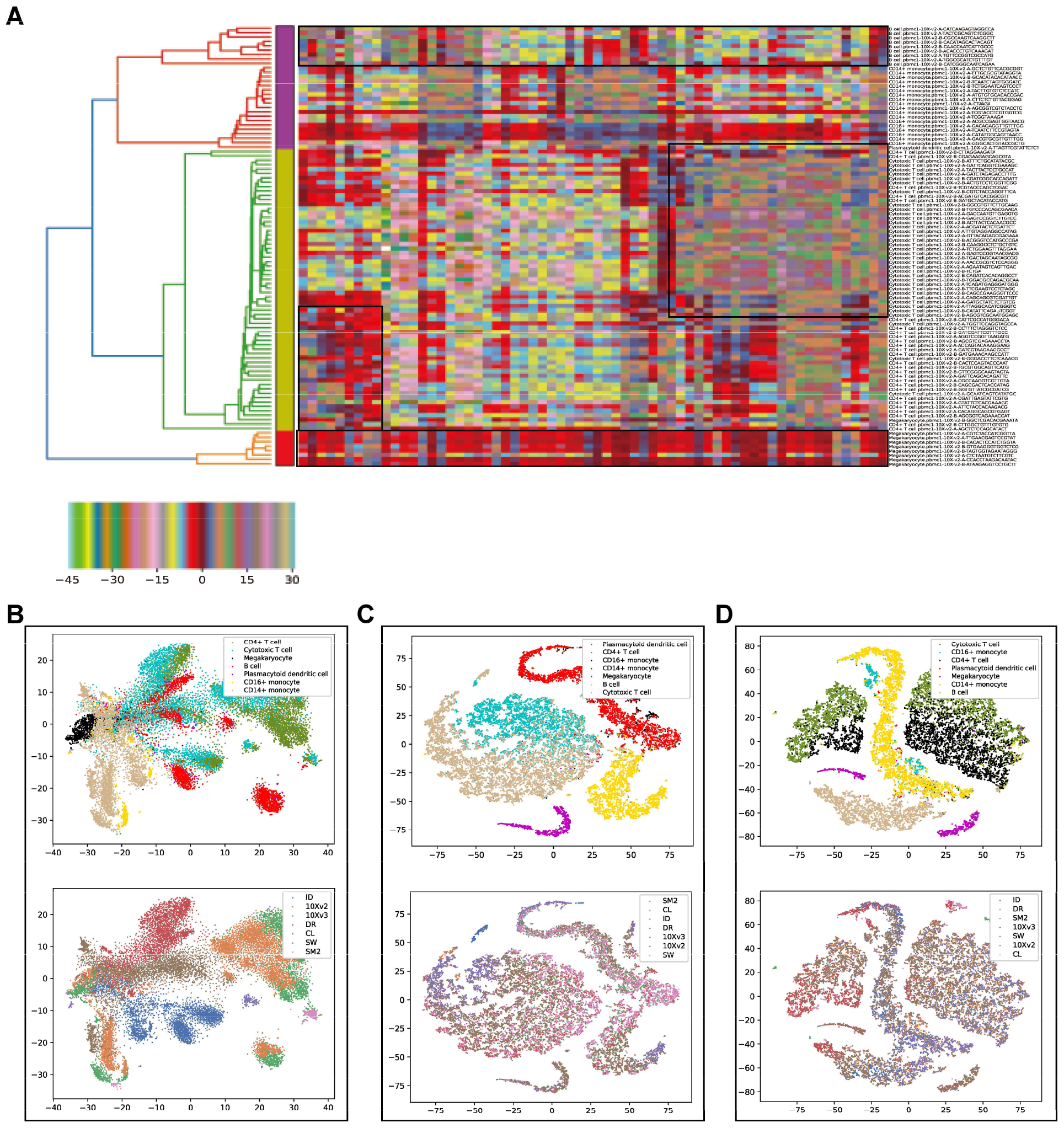
The analysis of distinguishing power of cell features extracted by scRCA. (A) Analysis of extracted CT features by running biclustering for 100 cells (covering 5 cell types) selected from the pbmc1_10Xv2 dataset. For each line of text, ‘.’ Before is the true cell type and after is the cell ID in the query dataset. (B) The t-SNE plots of the raw query dataset. (C) The t-SNE plots of CT features of the query dataset extracted by the trained network g using the pbmc1_ID with S-15% as the reference dataset. (D) The CT features of the query dataset extracted by the trained network g (the pbmc1_ID with P-15% is the reference dataset) were mapped into the 2D space using t-SNE. In subfigure (B-D), cells are colored by true cell type and sequencing protocol. Other results are provided in Supplementary Figs. S6 and S7.

As shown in **Fig. 4A**, the clustering tree aligns with the cell type hierarchical structure, except for several ‘CD4+T cells’. In addition, cells with identical cell types share similar features, while cells of different types show different features. Similarly, the clustering tree on the query dataset also aligns well with the cell type hierarchical structure, despite that cells were RNA-sequenced via different protocols (**Supplementary Fig. S3**). In contrast, a number of cells did not cluster well using the raw gene transcription data produced using either one or multiple protocols, as evidenced by **Supplementary Figs. S4-S5**. These results suggest that CT features calculated by scRCA models, even when trained using imperfect reference datasets, contribute to its cell type annotation performance.

### scRCA overcomes batch effects caused by different experimental protocols

In **supplementary Fig. 3**, cells that were RNA-sequenced using distinct protocols still clustered well, suggesting that scRCA has the potential to overcome batch effects caused by using different experimental protocols. To thoroughly investigate batch effects introduced by particular protocols, in this section, we projected all query cells into a two-dimensional (2D) space (**Fig. 4B-D**) using *t*-SNE [50]. In the upper panel of **Fig. 4B**, there is a significant batch effect for cells sequenced using different protocols, e.g., B cells were dispersed. However, in the lower panels of **Fig. 4C** and **Fig. 4D**, the batch effect induced was alleviated after being processed by scRCA. Particularly, the cells of the same type from different protocols clustered well. These results (**Fig. 4B-D** and **Supplementary Fig. S6-S7**) suggested that scRCA overcomes batch effects caused by different sequencing protocols.

### scRCA reveals marker genes for individual query cells

The interpreter module (‘Materials and Methods’) allows scRCA to discover the determining genes (section ‘Identifying potential marker genes for different cell types’) for each query cell. As shown in **Fig. 5A**, scRCA annotated a query cell as a ‘cytotoxic T cell’ based on the transcription values of multiple genes (i.e., NKG7 > 1.39, ECSIT <= 0.00, FBXW7 <= 0.00, MAPK14 <= 0.00, GNLY > 0.69, CCL5 > 1.95, and PPP2R5C > 0.69). Based on this information, NKG7, GNLY, CCL5, and PPP2R5C can be defined as the determining genes for cell type annotation. As such, the higher the transcription value for these genes, the more likely it is for the query cell to be annotated as ‘cytotoxic T cell’. We then sought experimental evidence for the annotation. By searching the CellMarker database [51], we identified and confirmed that NKG7, GNLY, and CCL5 are the marker genes of cytotoxic T cells. Similarly, as shown in **Fig. 5B**, IGKC, CD74, and CD79A were identified as the marker genes of the query cell (‘B cell’). However, as shown in **Fig. 5A** (bottom right), scRCA incorrectly annotated some cell types and the scRCA interpreter also indicates transcription levels of which genes are causing the prediction errors, thus alerting us to analyze these genes even further.

**Fig. 5.**
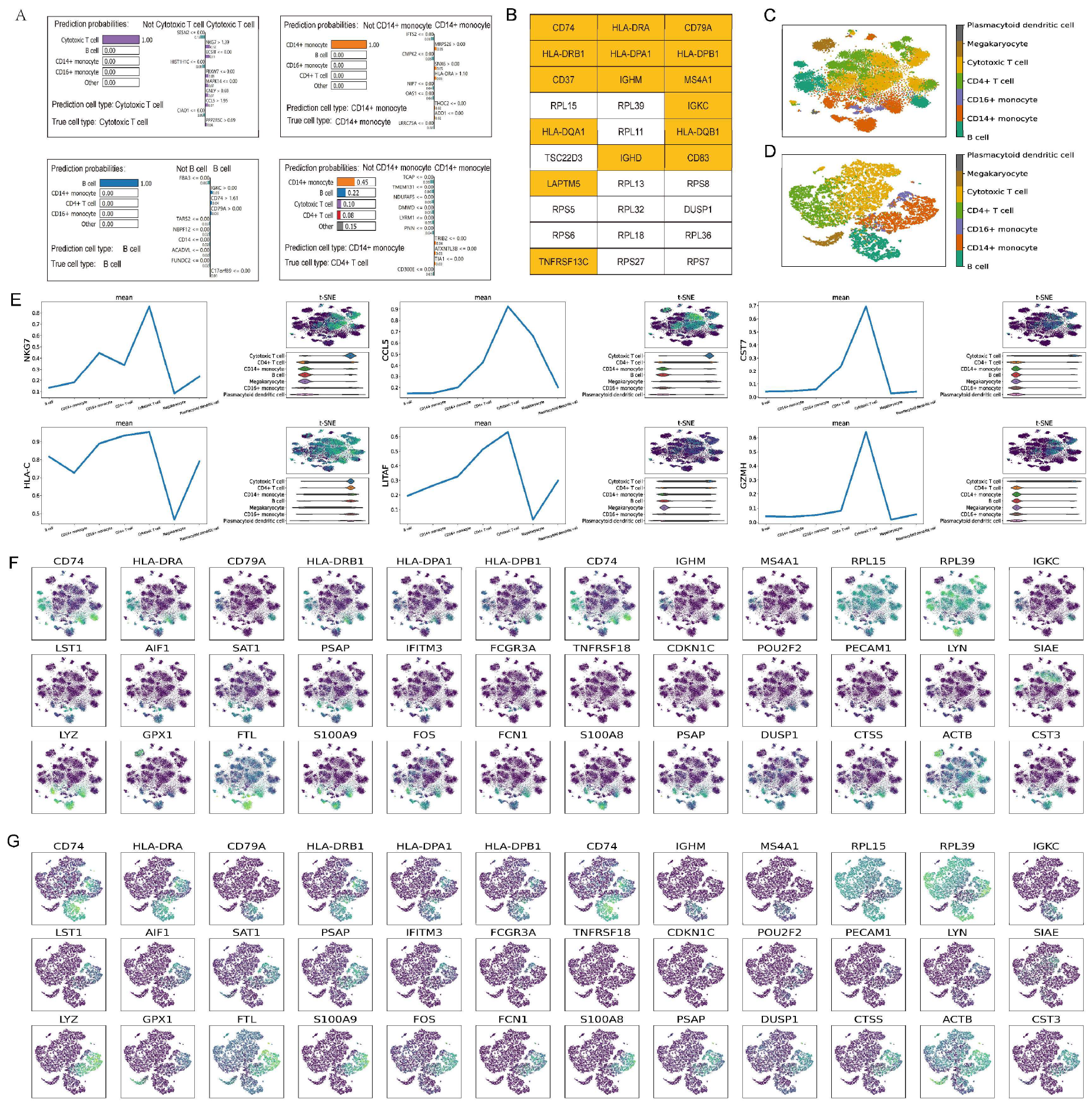
Uncovering the intrinsic mechanism of cellular annotation by scRCA. (A) The annotated results of scRCA. In each sub-figure, we provided the prediction probabilities, predicted cell type, factors (i.e., top 10 positive and negative gene transcription values), and the actual cell type. (B) The top 30 potential marker genes of B cells, for which genes are marked in yellow, are the confirmed marker genes; (C) The t-SNE plots of the query dataset (all genes projected by PCA using 16 principal components) using pbmc1_ID with S-15% as the reference dataset; (D) The t-SNE plots of the query dataset demonstrating the potential marker genes of various cell types projected by PCA using 16 principal components employing pbmc1_ID with S-15% as the reference dataset; (E) Plots showing the transcription of several potential marker genes of cytotoxic T cells, including the average expression value for individual cell types, gene expression value on t-SNE projection, and violin plots; (F) Each row represents the transcription levels of marker genes predicted by scRCA for B cells, CD16+ monocytes and CD14+ monocytes, respectively, overlaid on the all genes t-SNE plot of the query dataset; (G) Each row represents the transcription levels of marker genes predicted by scRCA for B cells, CD16+ monocytes and CD14+ monocytes, respectively, overlaid on subfigure D.

### scRCA infers marker genes for individual cell types

To investigate the marker genes for individual cell types, we re-ran scRCA using the corrupted pbmc1_ID dataset by symmetric flip using a noise level of 15% in the reference dataset to train networks *f* and *g*. The query dataset was generated by combining all other pbmc1 datasets and the pbmc2_ID dataset (‘Materials and Methods’). We then predicted marker genes (**Fig. 5B** and **Supplementary Fig. S8**) for individual cell types in the query dataset (refer to the ‘Identifying potential marker genes for different cell types’ section in ‘Materials and Methods’). scRCA discovered 16 marker genes confirmed by CellMaker for B cells in the potential maker gene list (**Fig. 5B**). It is important to note that other genes may be associated with its cell type but have not yet been investigated. In addition, according to **Fig. S8A-E**, scRCA can discover the confirmed marker gene(s) for CD14+Monocyte, CD16+Monocyte, Cytotoxic T cell, CD4+T cells, and Megakaryocyte cells. From **Supplementary Fig. S9**, the marker genes predicted were shown to be enriched for particular functions in various cell types. In **Fig. 5C-D**, all shared genes and potential marker genes were used to cluster cells represented in the query dataset, respectively. According to the results, the same cell types clustered better when marker genes were used. These results provide evidence that scRCA can overcome batch effects by selecting marker genes of each cell type in different datasets. In **Fig. 5E**, the average transcription values of genes CST7 and GZMH in cytotoxic T cells are substantially higher than in CD4+ T cells. Therefore, scRCA can find the key genes that distinguish the cell subtypes. The marker genes predicted here are clearly distinctive based on **Fig. 5F-G** and **Supplementary Fig. S10**. For instance, the CD74 gene was highly expressed in B cells and CD14+ monocytes, whereas the CD79A gene was highly expressed in B cells. These results indicate that scRCA can identify marker genes from reference datasets that are specific to particular cell types, and that query cells can be reliably annotated using these genes.

### scRCA can be used as a prominent tool for the discovery of disease-specific markers

We explored the use of scRCA to distinguish cancer cells from normal cells, based on the interpretability of scRCA to infer disease-specific marker genes. A single-cell dataset of peripheral blood immune cells from two healthy donors and four patients diagnosed with multiple myeloma [52], containing 35,159 cells with 32,527 genes, was used to validate our method. Overall, we obtained an average recognition accuracy of 99% using five-fold cross-validation. scRCA identified both known and novel gene markers for multiple myeloma within the peripheral blood immune cells (**Fig. 6C**). For example, published work has shown that the TPT1 gene, which is broadly upregulated in peripheral immune cells from multiple myeloma patients, is associated with multiple myeloma [53]. scRCA also identified that the S100 protein family (including S100A4, S100A6, S100A8, and S100A9) is associated with multiple myeloma – consistent with the previous studies [54]. Dysregulated expression of S100 protein family members is associated with cancer proliferation, invasion, angiogenesis, and inflammation. Other work suggested that the extracellular protein S100A9 promotes multiple myeloma and that the inhibition of S100A9 might have a therapeutic benefit [55]. In addition, a recent study showed that the S100A4 protein expression in multiple myeloma was associated with poor survival, and protein S100A8 and S100A9 were markers associated with a poor response to treatment in multiple myeloma patients with proteasome inhibitors (such as bortezomib, carfilzomib, and ixazomib) and the histone deacetylase inhibitor panobinostat [56]. Taken together, these results demonstrate that scRCA is capable of identifying both known and novel marker genes, which is of critical importance in understanding carcinogenesis and therapy development.

**Fig. 6.**
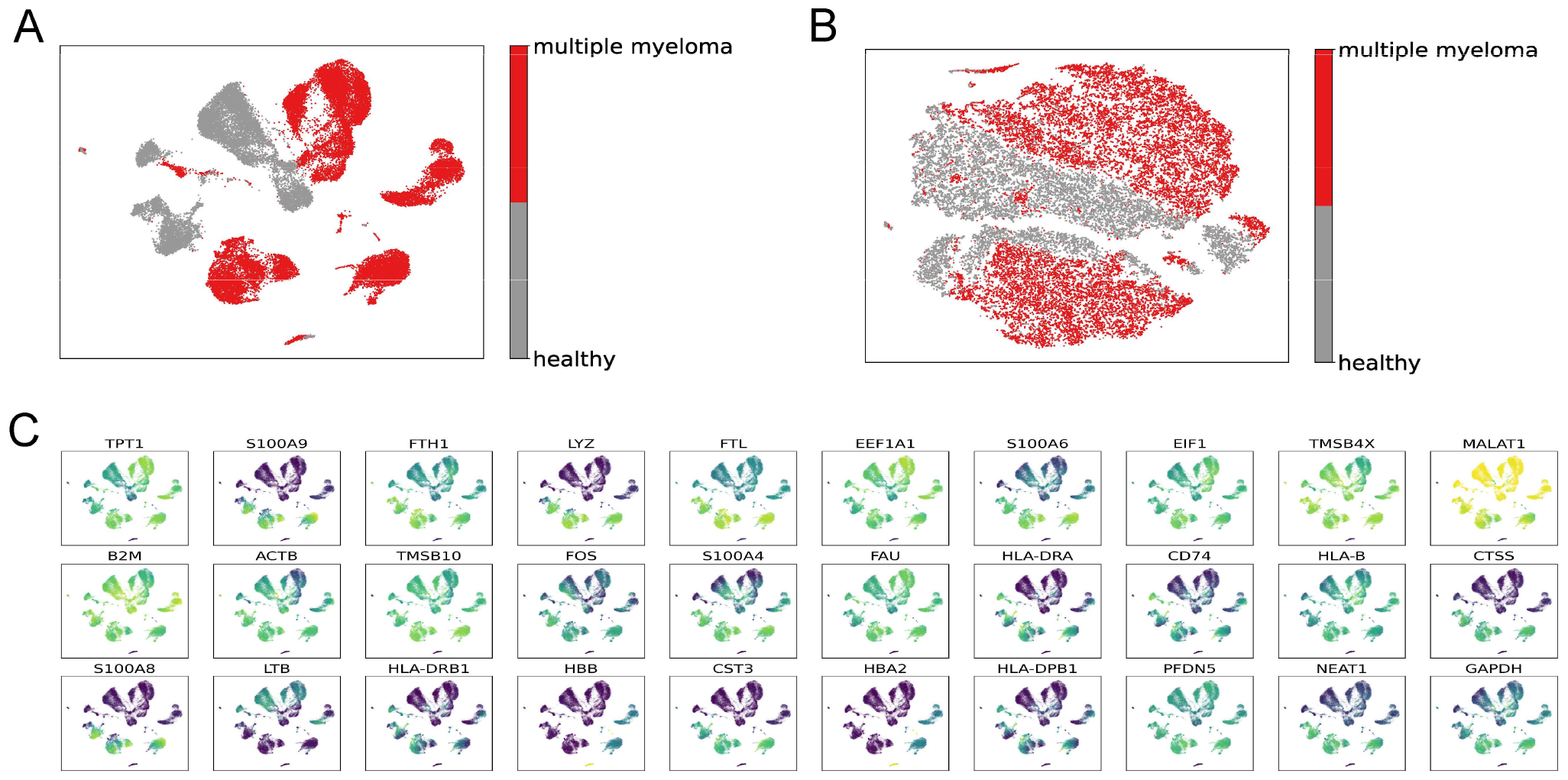
The discovery of disease-specific marker genes in the multiple myeloma dataset. (A) t-SNE plots of the entire filtered dataset. (B) t-SNE plots of the gene set selected using scRCA. (C) The expression level overlaid on t-SNE projection for disease-specific marker genes.

## Discussion

In this study, we established a novel deep learning-based approach, termed scRCA, to identify cell types using imperfect reference datasets. As annotation errors are inevitable in single-cell transcriptomic investigations, developing cell type annotation approaches that can extract relevant features from noisy reference datasets is critical. Despite some inaccuracies within the reference dataset, we have demonstrated that scRCA makes accurate cell type assignments. We implemented and compared the performance of three other cell type annotation methods based on different noise-robust loss functions, i.e., scFW, scGCE, and scDMI, and concluded that the accuracy of these methods improves significantly when pre-training is performed using the CCE loss function. In addition, we equipped a deep learning-based interpreter module within scRCA to increase the interpretability of annotations. Our benchmark experiments demonstrate superior annotation performance of scRCA compared to state-of-the-art reference dataset-based cell type annotators, including CaSTLe, scPred, SVM_rejection, Cell_Blast, mtSC, and DML. To explore the rationale for the annotation performance of scRCA, we showed that batch effects induced by distinct procedures can be mitigated and that the interpreter can discover known and novel marker genes for different cell types. Importantly, experiments conducted using multiple myeloma datasets showed that scRCA has the capacity to discover disease-associated markers.

Despite the outstanding performance of scRCA, there are some challenges that need to be addressed. For example, enabling scRCA to more accurately identify cell subtypes is an open challenge. Cell subtype classification is a fine-grained calssification problem that requires learning more detailed feature representations among cell subtype. The generalized large-margin (GLM) loss has been widely used in image fine-grained classification due to its ability to explore hierarchical label structure and similarity of fine-grained image categories. Therefore, GLM loss should ve considered to guild the network training in scRCA, thereby enabling cell subtype classification. In addition, scRCA cannot identify new cell types which are not represented in the reference dataset. In the future, we will endeavor to explore the usage of a metric loss function for in-depth cell subtype annotation. Furthermore, we will consider using a flexible threshold adaptation mechanism to scRCA for detecting novel cell types.

## Availability of data and materials

The datasets analyzed during the current study are publicly available. We download them from https://hemberg-lab.github.io/scRNA.seq.datasets/. An open-source implementation of the scRCA algorithm and code to reproduce the results can be downloaded from https://github.com/LMC0705/scRCA.

## Authors’ contributions

YL and GW conceived of Methodology, YL, HY and L-CS implemented the software and performed experiments. YL, CL, RBG, X-HH, J-NS, and D-JY all contributed to writing - review & editing. All authors read and approved the final manuscript.

