## Supplemental MATERIAL for "scRCA: a Siamese network-based pipeline for the annotation of cell types using imperfect single-cell RNA-seq reference data"

### Supplementary Text

#### Text S1

The scFW method is built on the forward (FW) loss function [1]. The FW loss is designed for loss correction, with the purpose of making it robust to the label noise. The FW loss is calculated as follows:

$$l_{\psi}\left(\tilde{Y}, \hbar(X)\right) = CCE\left(y, T^{T}\psi^{-1}\left(\hbar(X)\right)\right), \tag{1}$$

where $\psi$ is the link function (i.e., the softmax function in scFW), CCE() is the Cross-entropy loss, $\hbar(X)$ is the output of network part, $X$ is the gene expression of reference data, $\tilde{Y}$ is the incorrect cell annotation, and $T$ is the transfer matrix and can be approximated by two steps:

**Step 1:** Find the cell $x$ in the training data $X$ that is classified by scFW (herein, we used the CCE to pre-train the network part of scFW) as $y$ with the max probability, its mathematical expression is as follows:

$$\bar{x}^{i} = argmax_{x \in X}\, P\left(y = e^{i} | x\right). \tag{2}$$

**Step 2:** The transfer matrix can then be calculated as follows:

$$T_{ij} = p(y = e^{j} | \bar{x}^{i}). \tag{3}$$

**Text S2**

We set the network part of scDMI consistent with the scCCE, scFW, and scGCE but using the DMI loss function [2] to guide the training of the network. The DMI loss function is defined as follows:

$$L_{DMI} = -\log(DMI(\hbar(X), \tilde{Y})), \tag{3}$$

where $\hbar(X)$ is the output of network part, $X$ is the gene expression of reference data, $\tilde{Y}$ is the incorrect cell annotation, $DMI(\hbar(X), \tilde{Y})$ is the function determinant based mutual information of $\hbar(X)$ and $\tilde{Y}$ and can be calculated as follows:

$$DMI(\hbar(X), \tilde{Y}) = \left|det(Q_{\hbar(X),\tilde{Y}})\right|, \tag{4}$$

where $Q_{\hbar(X),\tilde{Y}}$ is the matrix format of the joint distribution $h(X)$ and $\tilde{Y}$ .

**Text S3**

The network framework part of scGCE is consistent with scCCE, scFW and scDMI using the GCE loss function [3] to guild the training of network part. The GCE loss is defined as follows:

$$\zeta_{GCE}\big(f(x), e_j\big) = \begin{cases} \zeta_q(k) & \text{if } f_j(x) \le k \\ \zeta_q\big(f(x), e_j\big) & \text{if } f_j(x) > k' \end{cases} \quad (5)$$

where $\zeta_q(f(x), e_j) = \frac{(1-f_j(x)^q)}{q}$, $q$ is the hyperparameter, ranges (0,1], and $k$ is a threshold. When the *f*-value (model output) is less than $k$, the sample is pruned and only the gradient of the cell samples with a *f*-value greater than $k$ are used to optimize the model. While the *f*-value of noisy data is often lower, using the threshold $k$ to prune can effectively avoid overfitting on noisy data. When training directly with this loss function, a potential problem is that at the beginning of the training phase, most of the softmax outputs may be significantly smaller than $k$, resulting in a sharp drop in the number of valid samples. In addition, pruning the samples based on the softmax values at the beginning of training is suboptimal. To address this problem, the GCE loss needs to be defined as follows:

$$\arg\min_{\theta}\sum_{i=1}^{n}\zeta_{GCE}(f(x_i;\theta), y_i) = \arg\min_{\theta}\sum_{i=1}^{n} v_i\zeta_q(f(x_i;\theta), y_i) + (1-v_i)\zeta_q(k), \quad (6)$$

when $f_{y_i}(x_i) > k, v_i = 1$, otherwise, $v_i = 0$, and $\theta$ is the parameters of the scGCE. Moreover, we can optimize the Eq. (6) as follows:

$$\arg\min_{\theta}\sum_{i=1}^{n} v_i\zeta_q(f(x_i;\theta), y_i) - v_i\zeta_q(k) = \arg\min_{\theta, w\in[0,1]^n}\sum_{i=1}^{n} v_i\zeta_q(f(x_i;\theta), y_i) - \zeta_q(k)\sum_{i=1}^{n} w_i\ . \quad (7)$$

Given any $\theta$, the $w$ is 1 if $\zeta_q(f(x_i\,;\theta), y_i) \le \zeta_q(k)$, and 0 otherwise. Finally, we used the Stochastic gradient descent (SGD) algorithm [REF] to optimize the network part of scGCE.

### Supplementary Tables and Figures

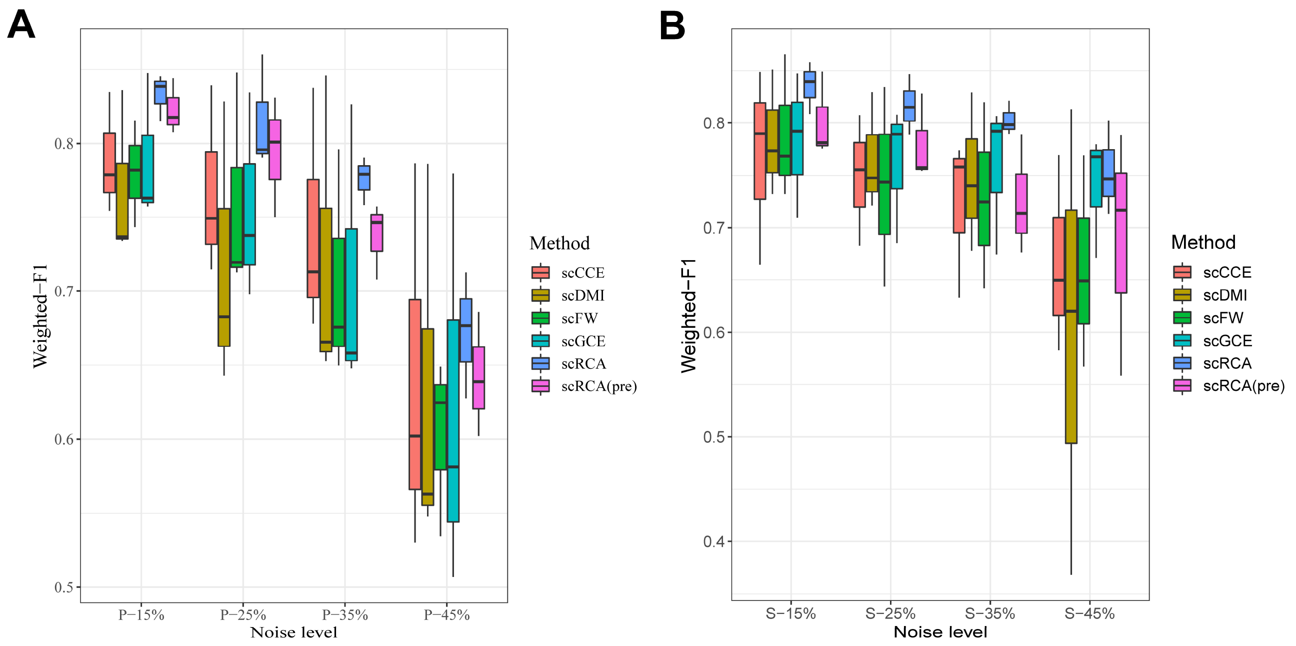


**Fig. S1.** The cell type annotation accuracy of each method using pbmc1_10Xv2, pbmc1_ID and pbmc1_10Xv3 as the reference datasets, under different cell type annotation corruption strategies and noise levels using (**A**) pair flip and (**B**) symmetric flip.

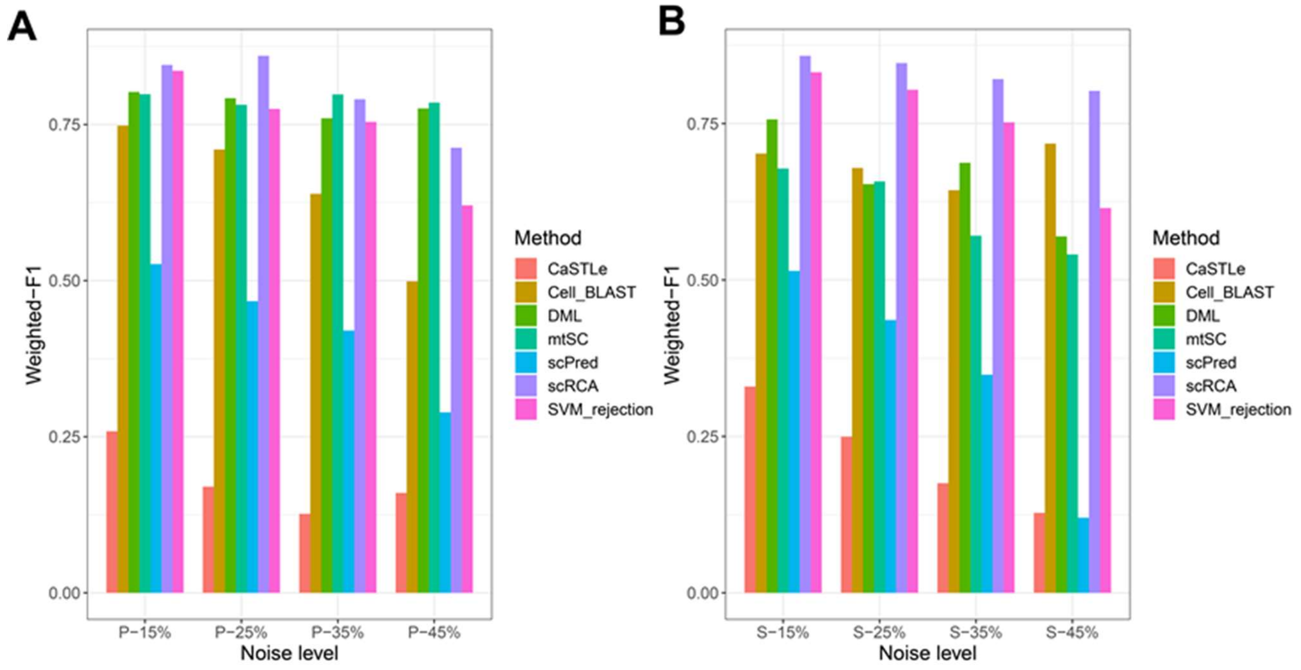


**Fig. S2.** Benchmarking (via Weighted-F1) scRCA and existing cell type annotation methods under different noise levels and cell type annotation corruption strategies: (**A**) pair flip and (**B**) symmetric flip using pbmc1_ID as the reference dataset.

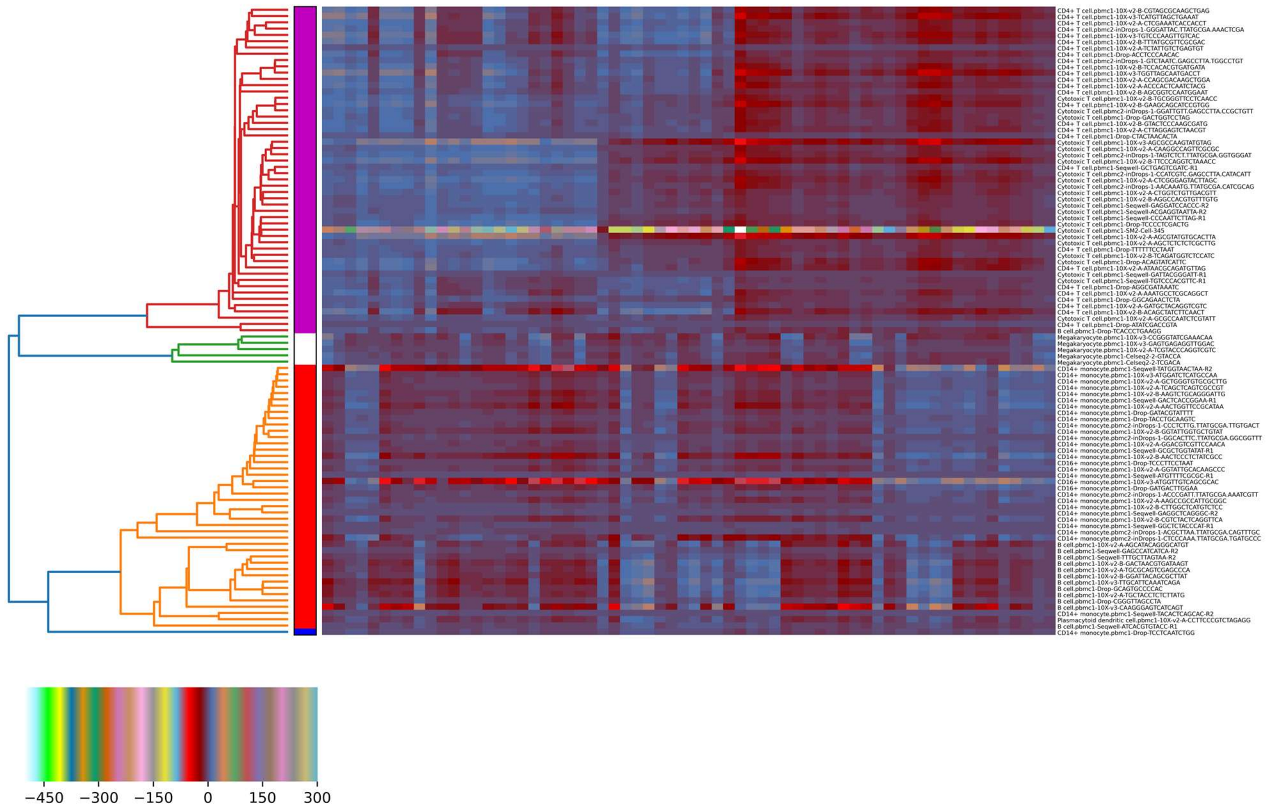


**Fig. S3.** Analysis of the extracted CT-features by running biclustering over 100 cells (covering 5 cell types) selected from datasets sequenced different protocols. Note that on the right side of the figure, cell types and names are presented before and after the "." for each cell, respectively.

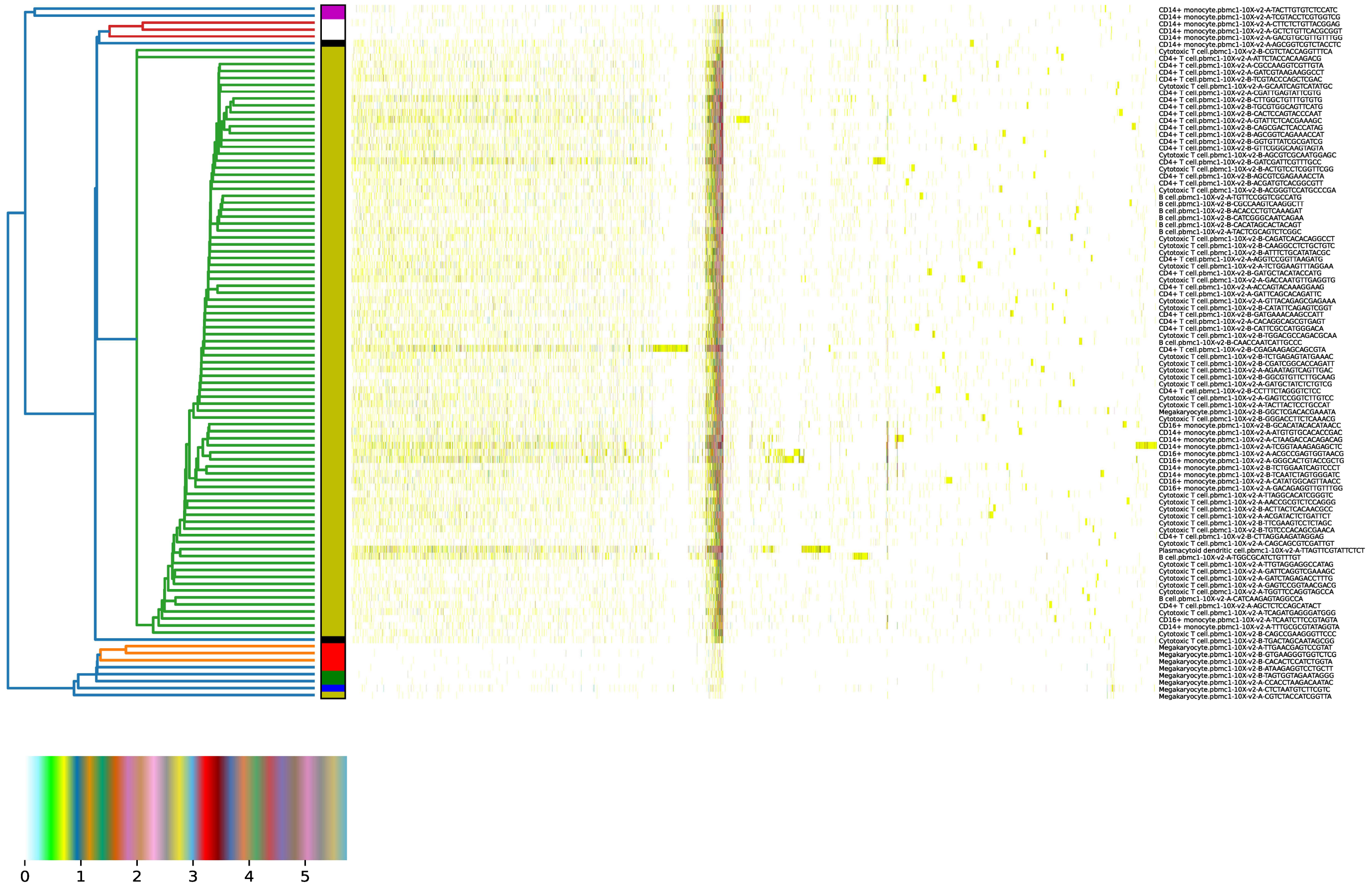


**Fig. S4.** Analysis of raw gene expression by running biclustering over 100 cells (covering 5 cell types) selected from the pbmc_10Xv2 dataset. On the right side of the figure, cell types and names are presented before and after the "." for each cell, respectively.

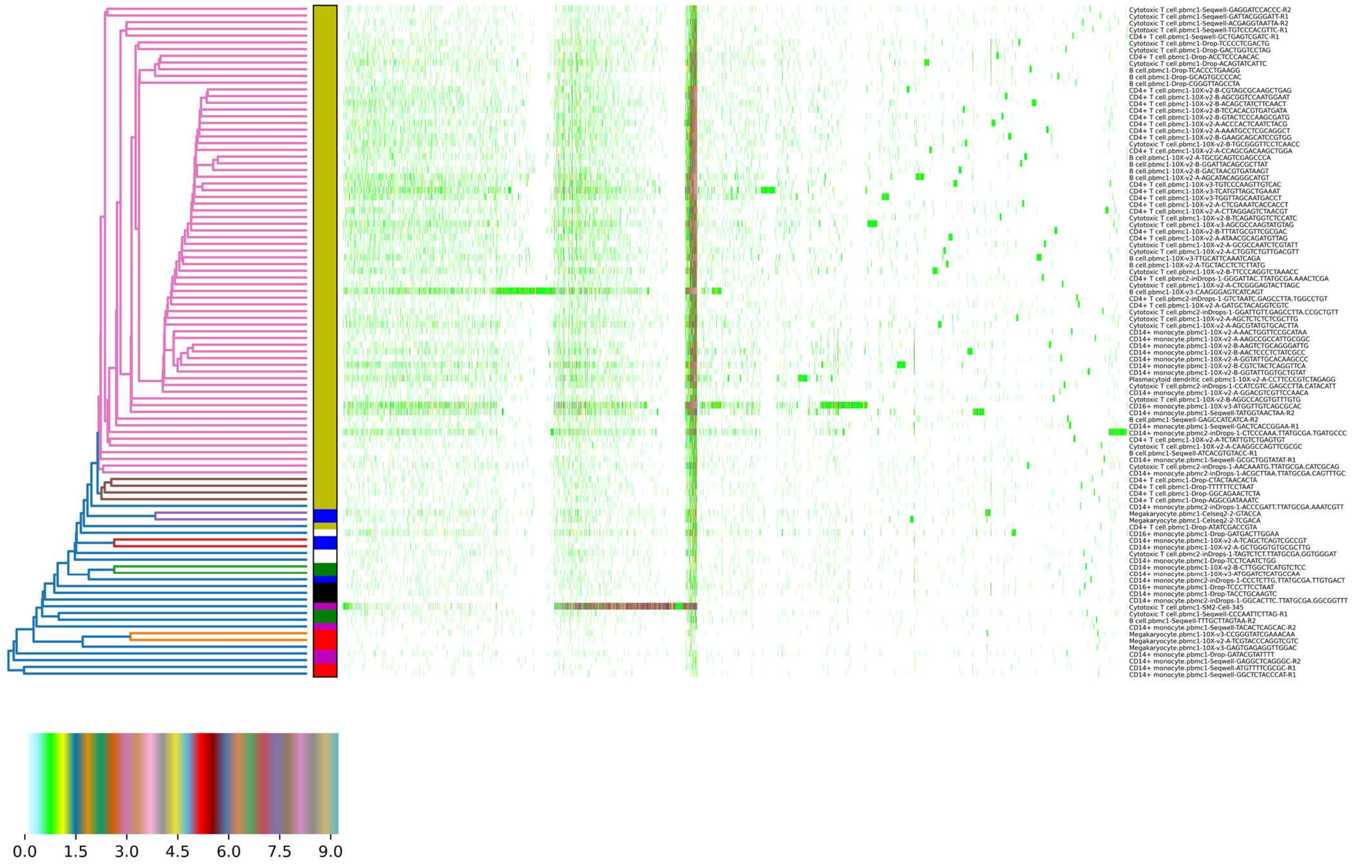


**Fig. S5.** Analysis of raw gene expression by running biclustering over 100 cells (covering 5 cell types) selected from datasets sequenced by different protocols. On the right side of the figure, cell types and names are presented before and after the "." for each cell, respectively.

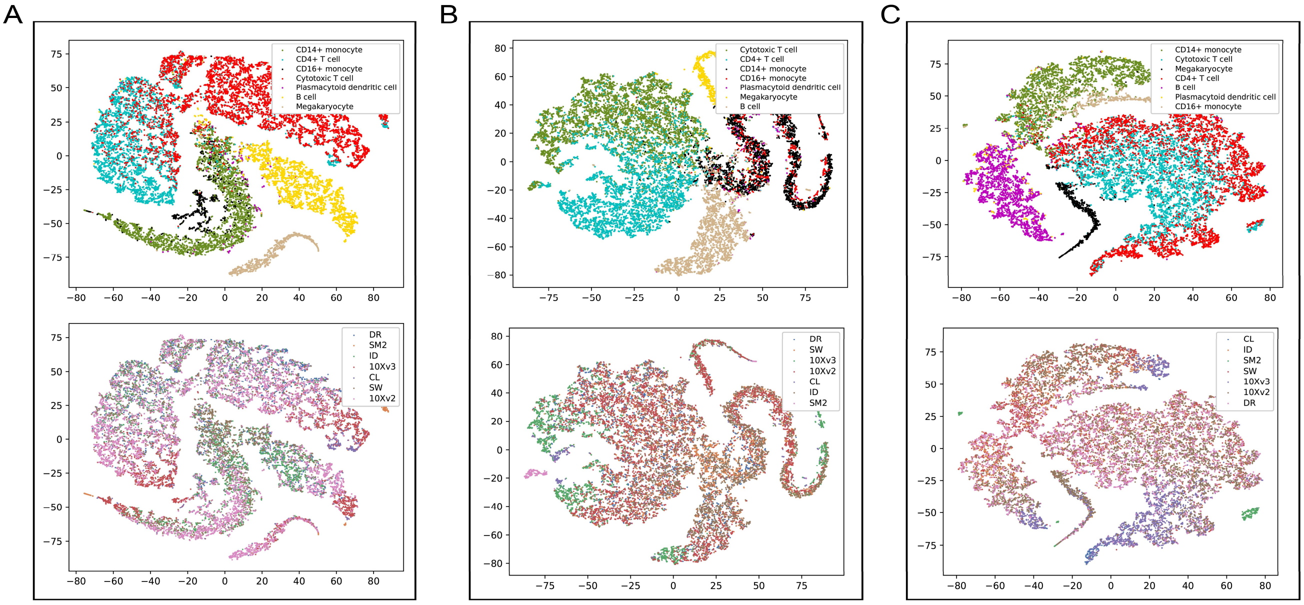


**Fig. S6. Qualitative evaluation of scRCA's ability to mitigate batch effects using *t*-SNE for visualization**. (**A**) The CT-features of test dataset extracted by trained network *g* were mapped into the 2D space using *t*-SNE using the pbmc1_ID with (**A**) pair flipping-25%, (**B**) pair flipping-35%, and (**C**) pair flipping-45%, as the reference dataset, respectively.

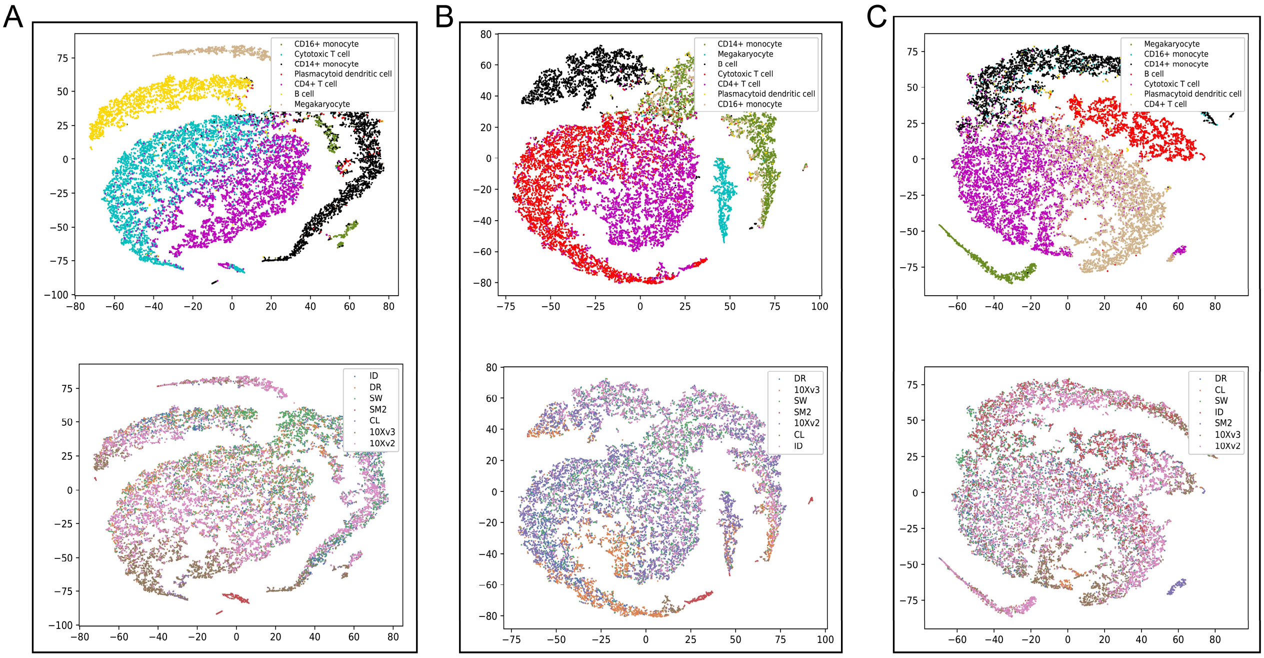


**Fig. S7. Qualitative evaluation of scRCA's ability to mitigate batch effects using *t*-SNE for visualization**. (**A**) The CT-features of test dataset extracted by trained network *g* were mapped into the 2D space using *t*-SNE using the pbmc1_ID with (**A**) sysmetric-25%, (**B**) sysmetric-35%, and (**C**) sysmetric-45%, as the reference dataset, respectively.

A

| | | |
|---|---|---|
| LYZ | GPX1 | FTL |
| S100A9 | FOS | FCN1 |
| S100A8 | PSAP | DUSP1 |
| CTSS | ACTB | CST3 |
| FCER1G | FTH1 | S100A6 |
| AIF1 | CSTA | VCAN |
| HLA-DRA | SAT1 | EMP3 |
| LST1 | GRN | TYMP |
| TKT | CD74 | AP1S2 |
| GAPDH | IFI6 | SERP1NA1 |

B

| | | |
|---|---|---|
| LST1 | AIF1 | SAT1 |
| PSAP | IFITM3 | FCGR3A |
| TNFRSF18 | CDKN1C | POU2F2 |
| PECAM1 | LYN | SIAE |
| SERPINA1 | CST3 | CTSS |
| C10ORF54 | COTL1 | NAP1L1 |
| FGR | SLC25A37 | TPM3 |
| TYROBP | FCER1G | CEBPB |
| CFP | LRRFIP1 | ATP2A2 |
| ZNF302 | ARPC5 | PLEK |

C

| | | |
|---|---|---|
| NKG7 | B2M | HLA-C |
| CCL5 | CST7 | LITAF |
| GNB2L1 | PPIA | GZMH |
| ACTB | IFITM2 | DAZAP2 |
| H3F3C | CD99 | CTSW |
| CALM1 | FLNA | PPP2R5C |
| TMSB4X | H3F3B | RPL32 |
| RPS15 | PFN1 | HLA-A |
| SH3BGRL3 | TPM3 | KLRD1 |
| PRF1 | SYNE2 | HLA-B |

D

| | | |
|---|---|---|
| PIK3IP1 | RPL13 | RPL21 |
| SARAF | RPS12 | LTB |
| RPL13A | RPS14 | RPS15A |
| RPS18 | RPL19 | RPSA |
| RPL34 | RPL7 | RPS8 |
| COX4I1 | LEPROTL1 | RPS29 |
| ARHGAP15 | RPL28 | FOXP1 |
| RPL22 | ZFP36L2 | RPS6 |
| RPS25 | JUNB | RPS4X |
| RPS28 | IL7R | RPL23A |

E

| | | |
|---|---|---|
| NRGN | HIST1H2AC | GPX1 |
| PPBP | PF4 | SDPR |
| GNG11 | CLU | TUBB1 |
| MAP3K7CL | PGRMC1 | TAGLN2 |
| MARCH2 | TUBA4A | NCOA4 |
| MMD | SPARC | CTSA |
| CALM3 | CST3 | RGS18 |
| FERMT3 | RGS10 | RNF11 |
| ACRBP | MAX | MYL9 |
| TPM4 | FKBP1A | OCT4 |

**Fig. S8. The top 30 positive genes of various cell types.** (**A-E**) genes marked in yellow are confirmed marker genes of CD14+ monocytes, CD16+ monocytes, Cytotoxic T cells, CD4+ T cells, and Megakaryocytes, respectively. The genes marked in blue in (**E**) are the marker genes of Natural killer T (NKT) cells.

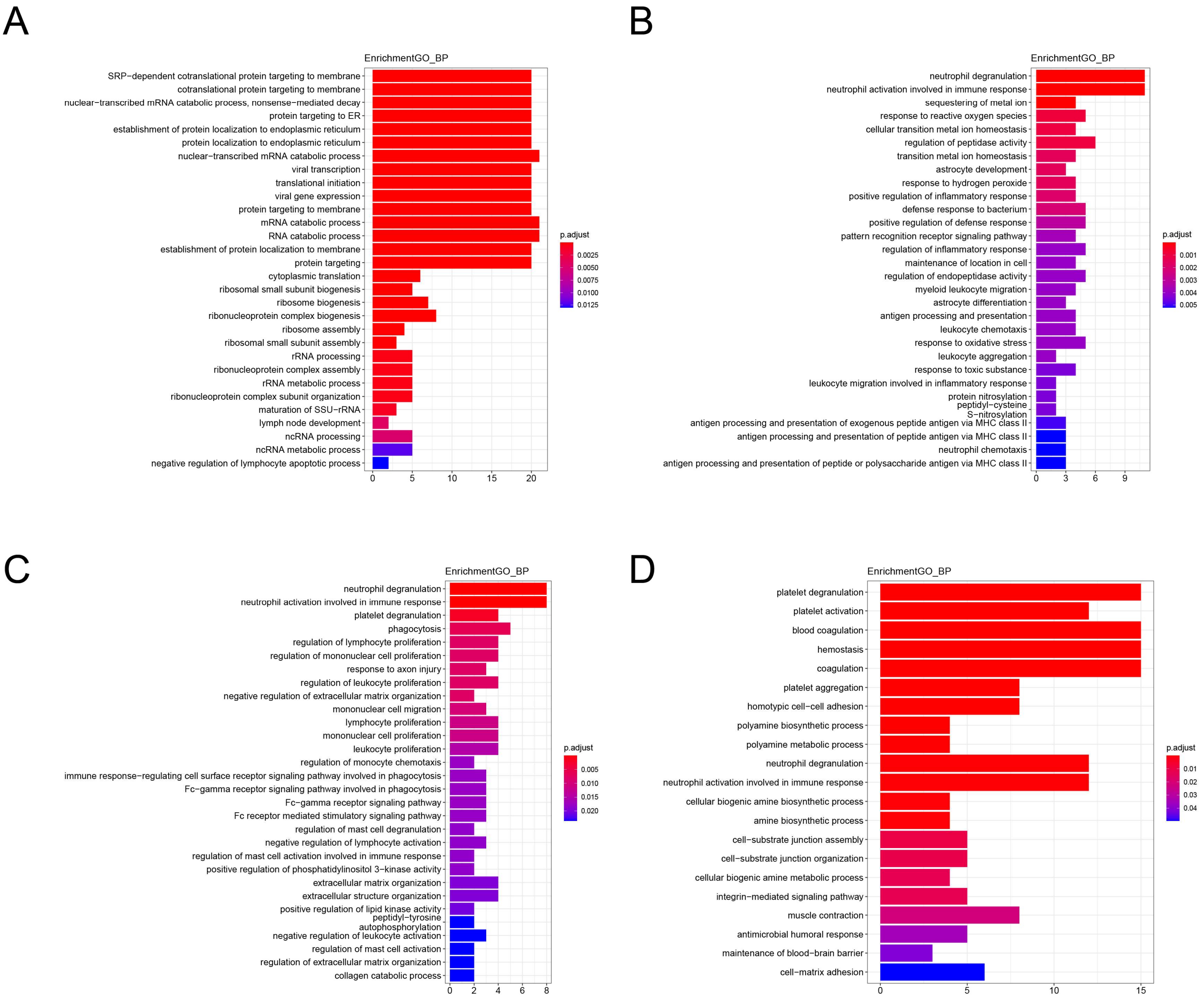


**Fig. S9. The Gene ontology (GO) enrichment analysis for the biological processes of the potential genes for different cell types:** (**A**) CD4+T cells; (**B**) CD14+ monocyte cells; (**C**) CD16+ monocytes; and (**D**) Megakaryocyte.

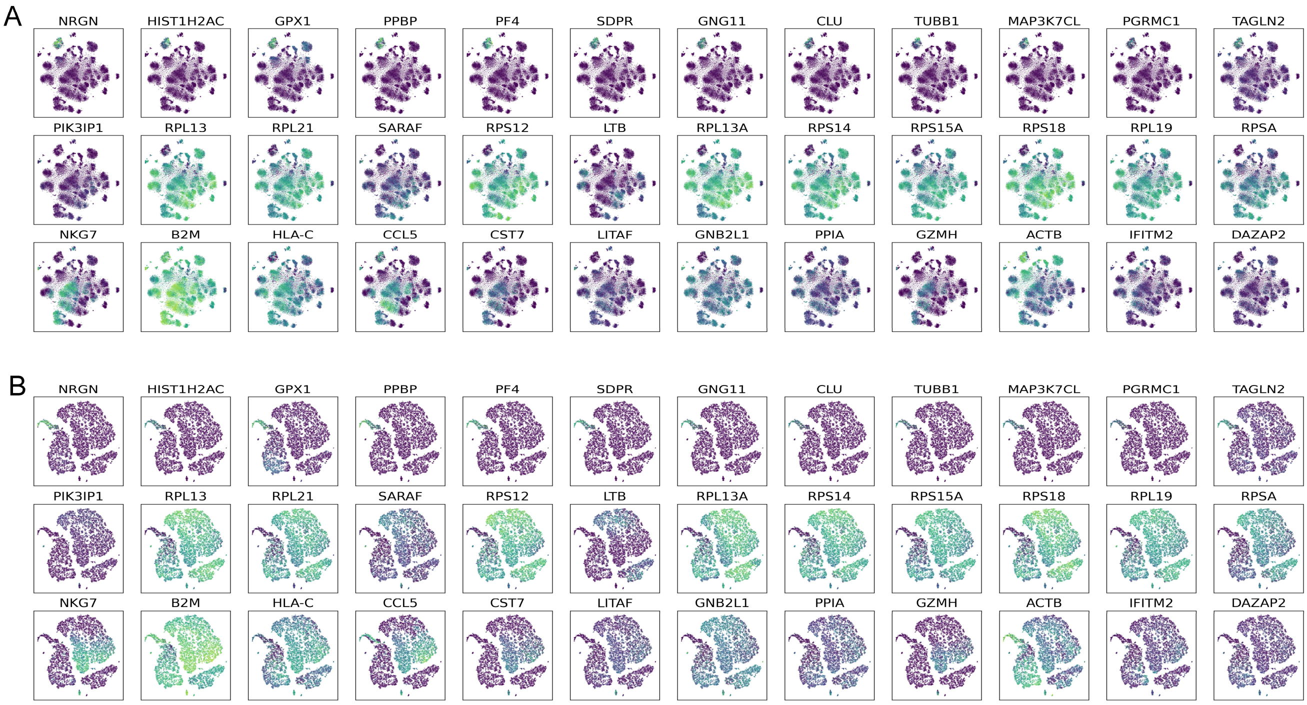


**Fig. S10. Expression level of additional selected genes overlaid on *t*-SNE plots.** (**A**) The expression level of the top 12 positive genes overlaid on the all genes *t*-SNE plot of the query dataset using pbmc1_ID with symmetric-15% as the reference dataset; (**B**) The expression level of the top 12 positive genes overlaid on the all positive trend genes t-SNE plot of the query dataset using pbmc1_ID with symmetric-15% as the reference dataset. The three columns of (A) and (B) are top positive genes for Megakaryocyte cells, CD4+T cellS and Cytotoxic T cells, respectively.

**Table S1.** An overview of the datasets used in this study

| Dataset | No. of cell Types | No. of genes | No. of cells | Protocol | reference |
|---|---|---|---|---|---|
| pbmc1_10Xv2 | 9 | 33,694 | 6444 | 10X version 2 | [4] |
| pbmc1_10Xv3 | 8 | 33,694 | 3222 | 10X version 3 | [4] |
| pbmc1_CL | 7 | 33,694 | 253 | CEL-Seq2 | [4] |
| pbmc1_DR | 9 | 33,694 | 3222 | Drop-Seq | [4] |
| pbmc1_iD | 7 | 33,694 | 3222 | inDrop | [4] |
| pbmc1_SM2 | 6 | 33,694 | 253 | SMART-Seq2 | [4] |
| pbmc1_SW | 7 | 33,694 | 3176 | Seq-Well | [4] |
| pbmc2_10Xv2 | 9 | 33,694 | 3362 | 10X version 2 | [4] |
| pbmc2_CL | 5 | 33,694 | 273 | CEL-Seq2 | [4] |
| pbmc2_DR | 6 | 33,694 | 3362 | Drop-Seq | [4] |
| pbmc2_iD | 9 | 33,694 | 3362 | inDrop | [4] |
| pbmc2_SM2 | 6 | 33,694 | 273 | SMART-Seq2 | [4] |
| pbmc2_SW | 4 | 33,694 | 551 | Seq-Well | [4] |
